# Reflection confocal microscopy for quantitative assessment of airway surface liquid dysregulation and pharmacological rescue in cystic fibrosis under near-physiological conditions

**DOI:** 10.1101/2024.03.05.583496

**Authors:** Ayca Seyhan Agircan, Marko Lampe, Heike Scheuermann, Tobias Albrecht, Simon Y. Graeber, Anita Balázs, Ingo Baumann, Stephan Block, Rainer Pepperkok, Marcus A. Mall, Julia Duerr

**Affiliations:** Department of Translational Pulmonology, University of Heidelberg, Heidelberg, Germany; Translational Lung Research Center Heidelberg (TLRC), Member of the German Center for Lung Research (DZL), Heidelberg, Germany; Advanced Light Microscopy Facility, European Molecular Biology Laboratory, Heidelberg, Germany; Department of Otorhinolaryngology, Head and Neck Surgery, University Medical Center of Eberhard-Karls University, Tübingen, Germany; Department of Otolaryngology, Head and Neck Surgery, Medical Center of the University of Heidelberg, Heidelberg, Germany; Department of Chemistry and Biochemistry, Freie Universität Berlin; Cell Biology and Biophysics Unit, European Molecular Biology Laboratory, Heidelberg, Germany; Department of Pediatric Respiratory Medicine, Immunology and Critical Care Medicine, Charité-Universitätsmedizin Berlin, Berlin, Germany; Berlin Institute of Health at Charité-Universitätsmedizin Berlin, Berlin, Germany; German Center for Lung Research (DZL), Associated Partner Site, Berlin, Germany

**Keywords:** Airway surface liquid, confocal reflection microscopy, cystic fibrosis, βENaC-Tg mice, airway epithelium

## Abstract

Proper regulation of airway surface liquid (ASL) is essential for effective mucociliary clearance (MCC) in healthy airways, and ASL depletion due to deficient cystic fibrosis transmembrane conductance regulator (CFTR)-mediated anion/fluid secretion plays an important role in the pathogenesis of mucociliary dysfunction and chronic muco-obstructive lung disease in patients with cystic fibrosis (CF). The current standard for quantitative measurements of ASL height is confocal fluorescence microscopy that has the disadvantage that it requires apical addition of volume for fluorescent staining, and hence perturbation of the ASL. Therefore, our aim was to develop a method that enables studies of ASL regulation under unperturbed conditions using reflected light by confocal microscopy of primary airway epithelial cultures grown at air-liquid interface (ALI). After apical volume addition to primary tracheal mouse cultures, confocal reflection microscopy yielded comparable ASL height as confocal fluorescence microscopy on cultures of wild-type mice, and was sensitive to detect ASL depletion on cultures of βENaC-Tg mice. Under unperturbed conditions, ASL determined by confocal reflection microscopy was significantly higher in wild-type and βENaC-Tg mice compared to values obtained by confocal fluorescence microscopy. Studies in normal and CF primary human airway epithelial cultures showed that confocal reflection microscopy was sensitive to detect effects of low temperature rescue and pharmacological modulation including improvement of CFTR function by VX-809 and VX-770 in cultures from CF patients with the F508del mutation. Our results support confocal reflection microscopy as a novel sensitive technique for quantitative studies of ASL regulation and response to therapeutic intervention under unperturbed near-physiological conditions in healthy and CF airways.

**NEW & NOTEWORTHY:** Measurement of airway surface liquid (ASL) height by confocal fluorescence microscopy is an important tool to investigate ASL dysregulation and effects of therapeutic strategies aiming at restoring ASL volume to improve mucociliary clearance and lung function in patients with cystic fibrosis. However, confocal fluorescence microscopy has the disadvantage that it requires apical addition of volume for fluorescent staining of the ASL leading to perturbation of its height and composition. Here, we developed confocal reflection microscopy as a new method that enables quantitative assessment of ASL on highly-differentiated primary airway epithelial cultures under unperturbed near-physiological conditions by detection of refracted light.

## INTRODUCTION

In the mammalian lung, the conducting airways are covered by a thin layer ( ̴7 µm) of airway surface liquid (ASL) that facilitates ciliary beating and is therefore essential for mucociliary clearance (MCC) of inhaled pathogens, allergens and other irritants (1). Proper regulation of ASL volume relies on coordinate secretion and absorption of salt and water by the airway epithelium. In this process, ion/fluid secretion is mediated by the cAMP-regulated Cl^-^ channel cystic fibrosis transmembrane conductance regulator (CFTR) in concert with the Ca^+^-activated and constitutively active Cl^-^ channels Anoctamin-1 (ANO1), also known as TMEM16A and solute carrier family 26 member 9 (SLC26A9), whereas absorption is mediated by the epithelial sodium channel ENaC (2-5). In patients with cystic fibrosis (CF), a spectrum of mutations in the *CFTR* gene leads to deficient CFTR-mediated Cl^-^/fluid secretion and enhanced ENaC-mediated Na^+^/fluid absorption resulting in ASL depletion and mucus hyperconcentration, which in turn impairs mucociliary clearance causing mucus obstruction, chronic bacterial infection, inflammation, and progressive structural lung damage (2, 3, 6-8). The recent development of small molecule CFTR modulators that improve folding, trafficking and gating and thereby restore function to the most common *F508del* mutation, but also a spectrum of rare CFTR mutations, enables a personalized approach to CF therapy (9-16). However, despite the established link between the underlying ion transport defect, ASL depletion and mucociliary dysfunction in the pathogenesis of CF lung disease, studies on the effects of CFTR modulators and other ion channel modulators on ASL regulation remain limited (14, 17-20). With a growing number of pharmacologic and genetic approaches becoming available to restore CFTR function in the airway epithelium, quantitative assessment of ASL dysregulation and response to therapy may facilitate preclinical development of novel drugs and precision medicine for individual patients with CF (8).

Several methods have been established to measure ASL height (14, 17, 21-24). The most common approach uses confocal fluorescence microscopy to measure ASL on highly differentiated primary airway epithelial cell cultures under air-liquid interface (ALI) conditions (14, 17, 18). However, this approach requires fluorescent labeling of the ASL and measurements outside the cell culture incubator at fixed time points limiting the temporal resolution of the assay and missing important information about kinetics of ASL regulation (21, 22) and the effective observation time is limited due to the endocytosis of the labeled dextran (21, 25). Further, fluorescent labeling of the ASL requires addition of volume to the surface of airway epithelial cultures leading to an acute perturbation of ASL height and composition. Therefore, measurements of ASL height with confocal fluorescence microscopy can only be conducted after reabsorption of the added volume, missing the possibility of unperturbed steady state measurements. Further, studies of the time course of ASL regulation in response to environmental stimuli or therapeutic intervention are hampered with this technique.

The aim of this study was, therefore, to develop a new method to study ASL height under unperturbed near-physiological conditions without the need of adding fluorescent labeling and additional volume to the ASL. To overcome these limitations of previous techniques, we developed confocal reflection microscopy using a widely available standard confocal microscope as a new approach to measure ASL height on highly differentiated primary airway epithelial cultures over long time periods without addition of labeling or volume to the ASL. First, confocal reflection microscopy was validated by comparison with the commonly used protocol of fluorescence confocal microscopy in tracheal epithelial cell cultures from wild-type mice. Second, confocal reflection microscopy was used to compare ASL height at steady state and ASL regulation upon apical volume challenge on tracheal cultures from wild-type mice and mice with airway-specific overexpression of ENaC (βENaC-Tg) (26). Finally, to determine its potential for quantitative assessment of therapeutic interventions, we used confocal reflection microscopy to determine effects of low temperature rescue (27°C) and ion channel modulators including the ENaC blocker benzamil, the Slc12a2 (NKCC1) inhibitor bumetanide and the CFTR modulator dual combination VX-809/VX-770 (lumacaftor/ivacaftor) on ASL height on primary human airway epithelial cultures from CF patients and non-CF controls.

## MATERIALS AND METHODS

### Experimental animals

All animal studies were approved by the local animal welfare authority (35-9185.81/G-97/14, Regierungspräsidium Karlsruhe, Germany). The generation of βENaC-Tg mice (RRID:MGI:5698383) has been described previously (26). Mice backcrossed onto a C57BL6/N background (27) and wild-type littermates served as controls. Tracheal tissues from mice for cell culture were collected at the age of 6 – 20 weeks. Mice were housed in a specific pathogen-free animal facility with free access to food and water.

### Primary murine tracheal epithelial cultures

For each individual experiment tracheae from 10 mice per group were freshly excised and pooled. Epithelial cells were isolated as previously described (28) and cultured under ALI conditions on transwell membranes (Transwell-COL, Corning GmbH, Kaiserslautern, Germany). Cell cultures were used for experiments about 14 days after seeding, when trans-epithelial electrical resistance (TEER) measurements showed cell confluence (TEER >200 Ω/cm^2^).

### Primary human airway epithelial cultures

Patients were recruited at the Department of Otolaryngology, Head and Neck Surgery of the University Hospital Heidelberg (Heidelberg, Germany). The study was approved by the Ethics Committee of the University of Heidelberg (S136/2016), and written informed consent was obtained from all subjects. Primary human airway epithelial cells were freshly isolated from nasal tissue obtained from patients undergoing polyps resection (CF patients, n=8) or correction of septum deviation (non-CF controls, n=7) and cultured under ALI conditions as previously described (29-31). Age and genotypes of study participants are listed in table 1.

**Table 1:** Demographics of non-CF and CF donors of primary airway epithelial cells.

|  | Non-CF controls | CF patients |
| --- | --- | --- |
| Age, years (mean ± SD) | 34 ± 14 | 20 ± 8 |
| Sex, n (females/males) | 4/3 | 4/4 |
| Pancreatic insufficiency, n (%) | Not determined | 8 (100 %) |
| *CFTR* genotype (n) | Not determined | F508del/F508del (4) |
|  |  | F508del/Q220X (1) |
|  |  | F508del/G542X (1) |
|  |  | F508del/2183delAA->G (1) |
|  |  | F508del/R553X (1) |

### Immunofluorescence microscopy

Differentiated cell monolayers were fixed with ice-cold methanol and permeabilized with 0.2% (w/v) Triton X-100. After blocking with 1% (w/v) bovine serum albumin, cells were incubated with polyclonal rabbit anti-βENaC (32) and monoclonal mouse anti-acetylated α-tubulin (Thermo Fisher Scientific Cat# 32-2700, RRID:AB_2533073 and Cat# A-11070, RRID:AB_2534114) primary antibodies. Cells were rinsed with PBS and further incubated with 1:200 dilution of respective Alexa Fluor labeled F(ab’)_2_ fragment (Thermo Fisher Scientific Cat# A-11017, RRID:AB_2534084) together with 1:2000 dilution of Hoechst (Abcam) to counterstain cell nuclei. Cells were mounted with FluorSave medium (Merck Millipore). Images were obtained by using a confocal laser scanning microscope (Leica TCS SP8, Leica Microsystems) with suitable settings for respective secondary antibody labels and were processed with the open-source imaging analysis software Fiji (33, 34).

### ASL height measurements with fluorescence microscopy

All imaging was conducted at 37°C and 5% CO_2_ using a microscope incubator (EMBL, Heidelberg, Germany). Primary tracheal epithelial cultures were washed with PBS, and 20 μl of PBS containing 2 mg/ml rhodamine dextran (10 kDa; Thermo Fisher Scientific) was added to the lumen to visualize the ASL layer. Images of the rhodamine-labeled ASL were acquired by confocal microscopy (Leica TCS SP8, Leica Microsystems). The height of the ASL was measured by averaging the heights obtained from xz scans of 15 predetermined positions on the culture as previously described (35). ASL height was measured 5 min following the addition of the rhodamine dextran and at designated time points over a period of 24 h in primary tracheal epithelial cultures from βENaC-Tg mice and wild-type littermates.

### ASL height measurements with reflection microscopy

All imaging was conducted at 37°C and 5% CO_2_ using a microscope incubator (EMBL, Heidelberg, Germany) enclosing the entire stage, nosepiece and upper part of the microscope. To ensure humidity under typical cell culture conditions (>98%), a custom-made humidity chamber for the transwells was developed. An anodized aluminum holder in the size of a typical 96-well plate with a water reservoir and a central position to mount a 35mm coverslip-/sample-holder was manufactured and closed by a humidity tight lid. Exchange of gas and brightfield microscopy was enabled by a 50mm lummox dish with its gas permeable, clear membrane (Sarstedt) placed in the center of the lid. Equipment of similar performance is commercially available. 30 minutes prior to the measurements, cells were stained basolaterally with 1ng/µL calcein-AM (Life Technologies) without affecting the ASL. Images were captured using a Leica PlanApo CS2 20X / NA 0.75 multi-immersion objective (Leica) with immersion liquid (Immersol W 2010, Zeiss or distilled water) and correction collar was set to ̴0µm coverslip thickness and then manually adapted to optimize signal intensity of the lower reflection signal of the transwell membrane. As a substitute for a glass coverslip, a fluorinated ethylene propylene (FEP) membrane in the same size (24 mm diameter) was used. FEP membrane has the same refractive index as water and therefore no optical correction for this element was necessary compared to glass coverslips. The sample was placed on the FEP membrane on top of 2 pieces of parafilm (2mm width, 10mm length) on opposite sides to leave sufficient space for growth medium under the membrane. The reflection signal that is obtained due to the changes in the refractive index was recorded in XZ-scans using a 488 nm laser and a photomultiplier tube as detector (Fig. 1a). For long term experiments, samples were kept in the microscope for the time period and images were taken every 15 minutes at 15 defined positions in the middle of the well avoiding the edges. Image analysis were performed with the open-source imaging analysis software Fiji (33, 34).

**Figure 1.**
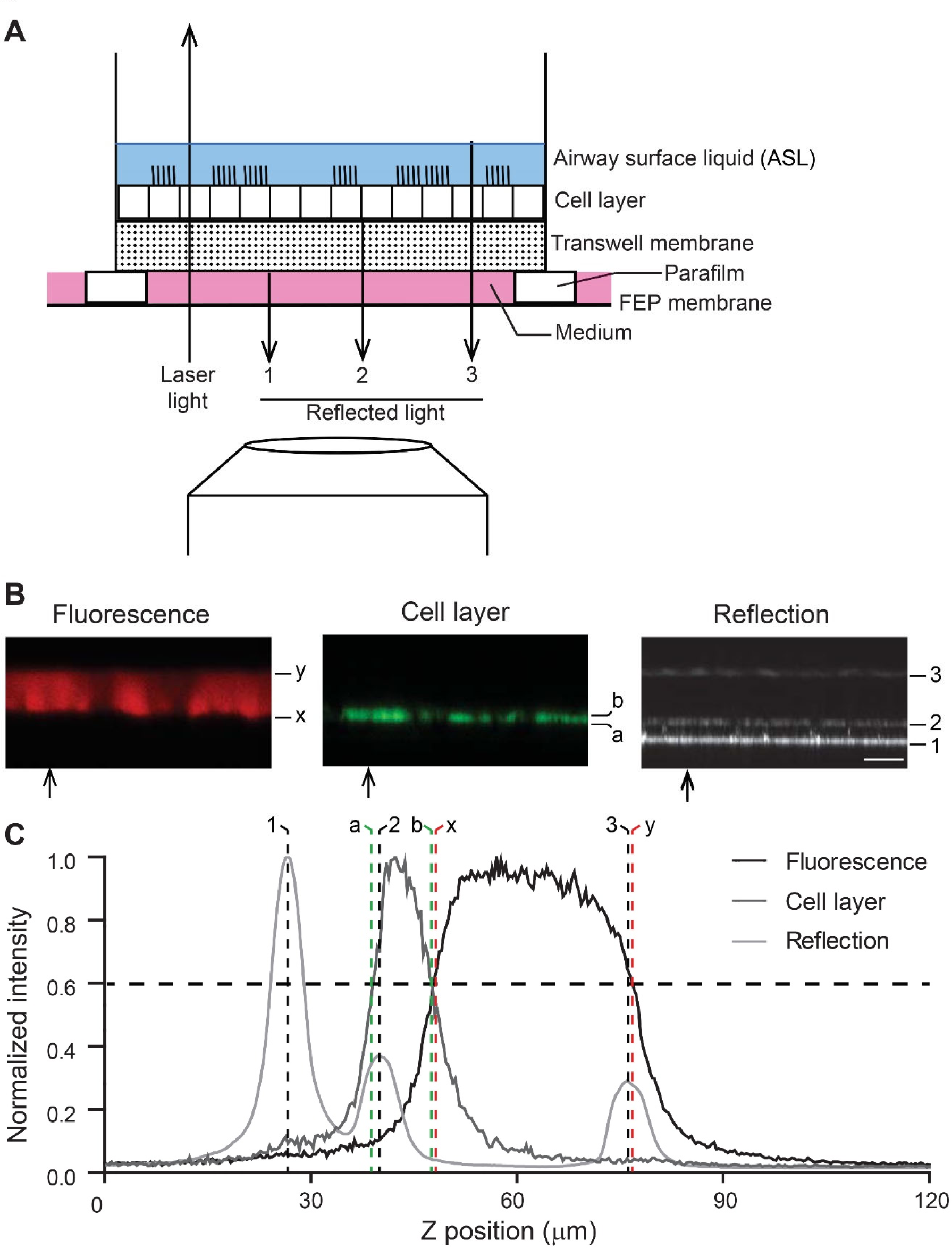
Principle of reflection confocal microscopy for measurements of ASL height on airway epithelial cultures. *A*: Schematic representation of the laser beam passing through a transwell with a differentiated airway epithelial layer grown at air liquid interface and the portions of light being reflected at each interface with a change in refractive index reversing its direction of propagation. For clarity, the reflected signal is depicted separately from the laser light. FEP: fluorinated ethylene propylene. *B*: Reflection signals obtained from normal (wild-type) murine primary tracheal epithelial cultures with the 488 nm laser by XZ-scanning and fluorescence images recorded in parallel at 488 nm for the cell layer (calcein-AM) and 561 nm for ASL (rhodamine dextran). Arrows mark the position at which the following line profiles of fluorescent intensities were taken from. Peaks of reflected light in the line profile are labeled correspondingly. In the fluorescence measurements (x) and (y) mark the basolateral and apical border of the rhodamine signal, respectively. (a) and (b) mark the borders of the calcein-AM signal representing the cell layer. For reflection measurement laser light is reflected at (1) the transition from medium to transwell, (2) the transition from transwell to cells, (3) the transition from ASL to air. Scale bar: 25 µm. *C*: Line profiles of an XZ-scan of the recorded reflection signal and the fluorescent signal from calcein-AM and rhodamine dextran. Vertical dashed lines mark the corresponding positions in the confocal images in *(B)*.

### Low temperature correction of F508del-CFTR

For low temperature correction of F508del-CFTR (36) in primary human airway epithelial cells derived from CF patients, cells were cultured under ALI condition at 37°C for 12 days and 48 hours prior to the ASL height measurements temperature was reduced to 27°C.

### Pharmacological treatment

To measure the effect of pharmacological modulation of ion transport in primary human airway epithelial cell cultures, baseline ASL height was measured and benzamil (100µM) was added to inhibit electrogenic ENaC-mediated Na^+^ absorption. Then 3-isobutyl-1-methylxanthine (IBMX; 100µM) and forskolin (1µM) were added to induce cyclic adenosine 5’-monophosphate (cAMP) mediated Cl^-^ secretion and bumetanide (100µM), an inhibitor of the basolateral Na^+^–K^+^– 2Cl^-^ cotransporter (NKCC1), was added to block transepithelial Cl^-^ secretion. The aforementioned substances were added basolaterally to the cell cultures. ASL height was measured continuously by taking images at every 15 minutes and for 45 minutes after addition of each compound. To determine effects of the CFTR modulator dual combination VX-809/VX-770 (lumacaftor/ivacaftor, primary human airway epithelial cells from CF patients with at least one F508del allele were cultured under ALI condition and incubated with 5µM VX-809 (Selleck Chemicals) basolaterally for 48 hours prior to ASL height measurements. Subsequently, 5µM VX-770 (Selleck Chemicals) was added acutely to the basolateral side as previously described (37) and continuous ASL height measurements were performed. All chemicals were obtained from Sigma-Aldrich if not stated otherwise and of the highest grade of purity available.

### Statistical analysis

Data were analyzed with GraphPad Prism software (GraphPad Software) and presented as mean ± SEM. Statistical analyses were performed using paired and unpaired, two tailed Student’s t-test or Wilcoxon signed rank test as appropriate, and p<0.05 was accepted as statistically significant. All data were obtained from at least three independent experiments.

## RESULTS

### Measurement of ASL height using confocal reflection microscopy

For initial testing of the feasibility of confocal reflection microscopy for quantitative assessment of ASL height, we first compared its performance to the established confocal fluorescence microscopy that requires fluorescent labeling of the ASL leading to an acute volume challenge on the apical surface of airway epithelia. These experiments were performed with primary tracheal epithelial cultures from wild-type mice grown under ALI conditions and signals detected by confocal reflection microscopy and fluorescence microscopy were recorded simultaneously in the same experiment (Fig. 1). The change in fluorescence intensity indicating the ASL height is flanked by signals of recorded reflected light that is obtained due to the changes in the refractive index and corresponding to the interface between upper transwell membrane and the basolateral membrane of the epithelial cells, and the interface between the ASL and the air at the surface of the ASL. To unambiguously distinguish epithelial cells from ASL under all experimental conditions, cells were labeled with calcein-AM added to the basolateral compartment, i.e. leaving the ASL unaffected. To determine ASL height by fluorescence, the half maximum intensities were used to determine the boundaries of the fluorescently stained liquid (distance between x and y; Fig. 1C). For reflection microscopy the distance between the peaks indicating the transitions between the transwell membrane and epithelial cells, and between ASL and air (number 2 and 3; Fig. 1C) was measured and cell height (distance between a and b; Fig. 1C) was subtracted to determine the ASL height on airway epithelial cultures. The overlay of spatial distribution of recorded change in fluorescence intensity and change in intensity of reflected light along the z-axis of scanned cultures at t=0 showed that measurements with both methods were equivalent (Fig. 1C).

### ASL height measurements with confocal reflection microscopy agrees with confocal fluorescence microscopy

To further validate confocal reflection microscopy for studies of ASL regulation over time, we next compared it to a standard protocol using conventional confocal fluorescence microscopy in primary tracheal cultures from wild-type mice for longitudinal measurements over a time course of 24 hours. Measurements started immediately after addition of rhodamine-dextran to the ASL. Fluorescence and reflection signals were recorded in parallel and the overlay of images showed exact matching of the distinct signals obtained with the two methods (Fig. 2A) producing the same values for ASL height at each time point over 24 hours (Fig. 2B). Taken together, under the same experimental settings, ASL height measurement by confocal reflection microscopy reliably reproduced ASL height measurement by confocal fluorescence microscopy including ASL regulation following the acute volume challenge required for fluorescence based measurements.

**Figure 2.**
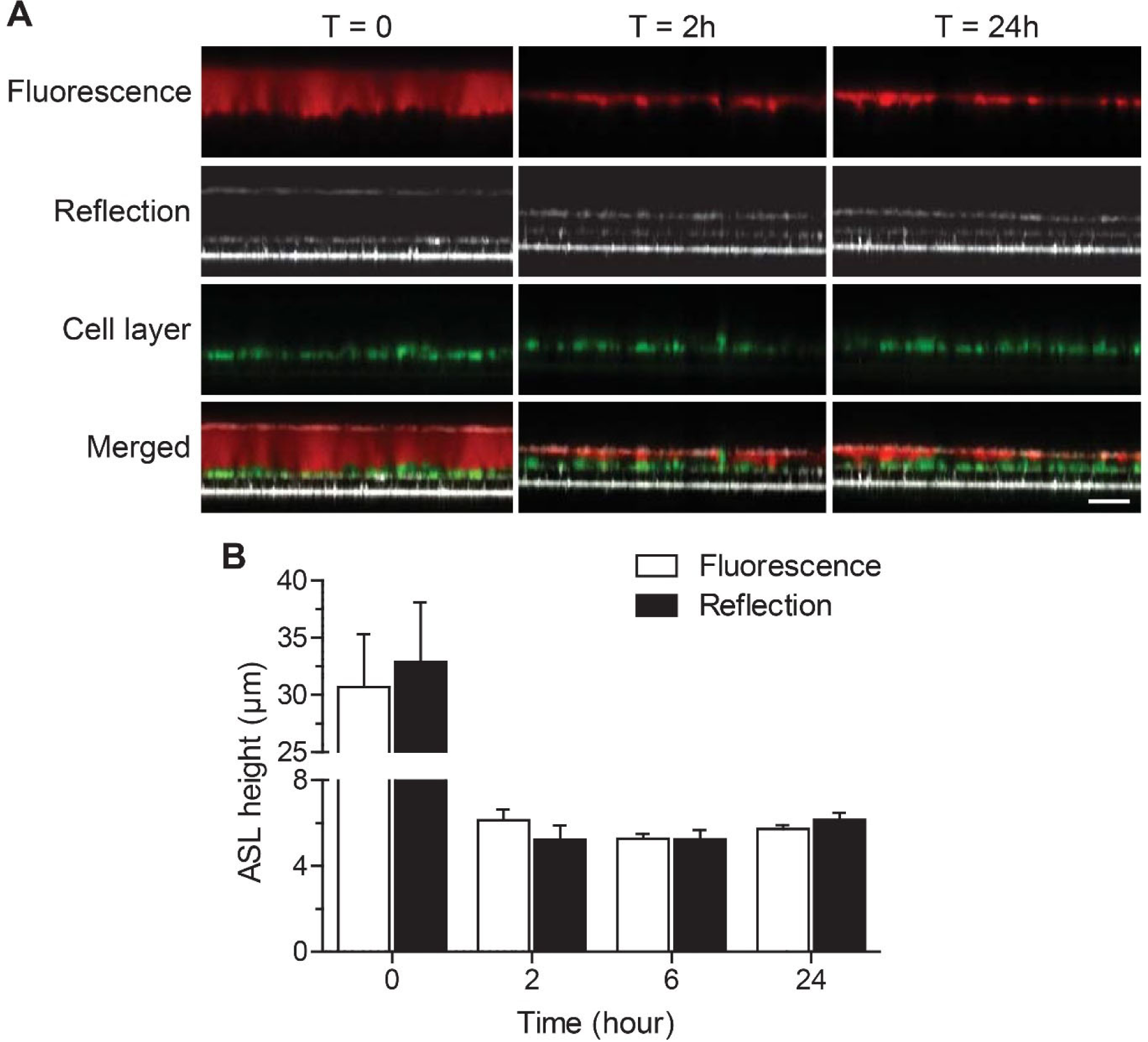
Comparison of ASL height after apical volume challenge of primary murine airway epithelial cultures determined by reflection vs. fluorescence confocal microscopy. Primary tracheal epithelial cell cultures from wild-type mice were grown at air-liquid interface for 14 days and 20 µl of fluorescent dye (rhodamine dextrane in PBS) was added to the apical compartment to label the ASL. A: Representative confocal images of the reflection signal recorded with the 488 nm laser by XZ-scanning and fluorescence images recorded in parallel at 488 nm for the cell layer (calcein-AM) and 561 nm for the ASL (rhodamine dextran). Images show ASL and labeled cells immediately after volume challenge (T = 0), 2 hours (T = 2) and 24 hours (T = 24) thereafter. Scale bar: 25 μm. B: Summary of ASL height as determined by confocal fluorescence microscopy and reflection microscopy immediately after dye addition, at 2 hours, 6 hours and 24 hours. n = 4 cultures per group.

### Confocal reflection microscopy detects ASL depletion in βENaC-Tg mice

To determine whether confocal reflection microscopy is sensitive to detect CF-like ASL depletion, we used this technique to compare ASL height on primary airway epithelial cell cultures from wild-type mice and βENaC-Tg mice that exhibit airway-specific overexpression of the β-subunit of ENaC, leading to increased epithelial Na^+^ absorption, ASL depletion, impaired MCC and CF-like lung disease (26, 38). Primary tracheal epithelial cells from mice of both genotypes were freshly isolated and cultured under ALI conditions resulting in a polarized epithelial cell layer as shown by immunofluorescence staining of acetylated α-tubulin as a marker for ciliated cells (Fig. 3A). Endogenous expression and overexpression of the βENaC-transgene were verified by immunofluorescence staining for βENaC and co-localization studies showed that wild-type and transgenic βENaC was expressed in non-ciliated cells (Fig. 3A). Measurements of numeric cell densities revealed that tracheal cultures from wild-type and βENaC-Tg mice consisted of ̴50% of ciliated cells similar to what has been reported for mouse trachea in vivo (39, 40) (Fig. 3B). In wild-type cultures, about 30% of all cells stained positive for βENaC, but no cells with high intensity βENaC staining were detected (Fig. 3AB). In cultures from βENaC-Tg mice, about 45% of total cells were βENaC positive and 20% of total cells, i.e. about the half of non-ciliated cells, showed high intensity of fluorescence when stained for βENaC confirming overexpression of βENaC under culture conditions (Fig. 3B).

**Figure 3.**
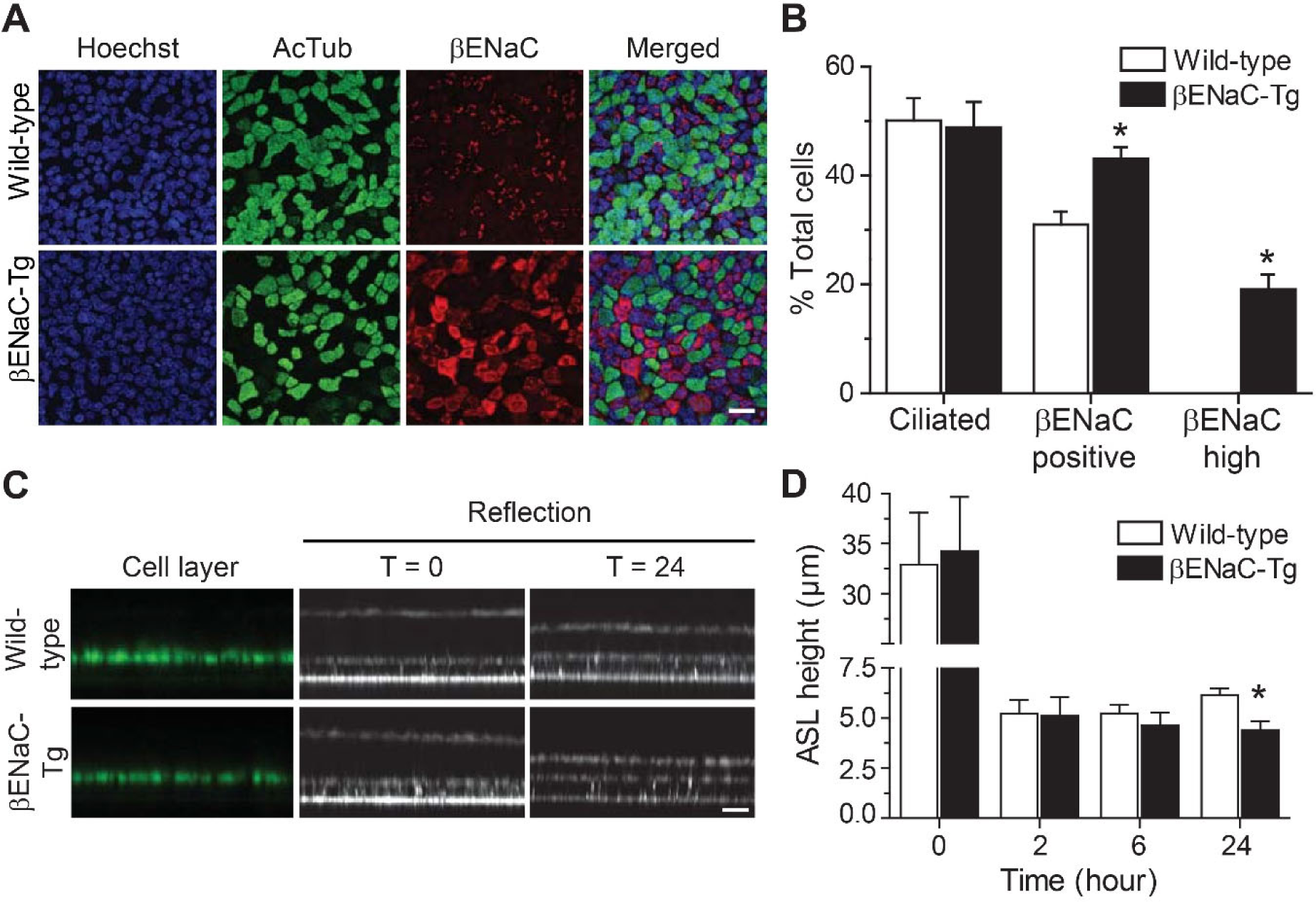
Reflection confocal microscopy detects steady state ASL depletion on primary airway cultures from βENaC-Tg mice after apical volume challenge. Primary tracheal epithelial cultures from βENaC-Tg mice and wild-type controls were grown at air-liquid interface for 14 days and numeric densities of ciliated cells, βENaC-expressing cells and regulation of steady state ASL height following an apical volume challenge (20 µl PBS) were determined. A: Representative images of wild-type and βENaC-Tg cultures co-immunostained with Hoechst (blue) as nuclear stain, anti-acetylated tubulin antibody (AcTub, green) and anti-βENaC antibody (red). Scale bar: 20 µm. B: Numeric cell densities of ciliated cells (AcTub-positive), βENaC-positive cells and βENaC-overexpressing cells were determined and expressed as percentages of total cells. n = 5 cultures per group, \**P* < 0.01. C: Representative confocal images of the reflection signal recorded with the 488 nm laser by XZ-scanning and calcein-AM staining showing the epithelial cell layer and ASL height immediately after apical volume challenge (T = 0) and 24 hours (T = 24) thereafter. Scale bar: 12 μm. D: Summary of ASL height as determined by reflection confocal microscopy immediately after apical volume challenge and at 2 hours, 6 hours and 24 hours thereafter. n = 4, \**P* < 0.01.

In these experiments, ASL height measurements by confocal reflection microscopy were performed following apical addition of fluorescent dye and volume for comparability with results of previous measurements performed with conventional confocal fluorescence microscopy (35, 38). Measurements started immediately after addition of rhodamine-dextran to the ASL. Fluorescence and reflection signals were recorded in parallel at 15 different predetermined positions of the cell culture well (Fig 3C and data not shown). After volume addition excess liquid was rapidly absorbed by cultures from wild-type and βENaC-Tg mice (Fig 3D). At 24 hours, when steady state was reached, ASL height determined by confocal reflection microscopy on βENaC-Tg cultures (4.4 ± 1.4 µm) was significantly reduced compared to wild-type cultures (6.1 ± 1.2 µm; *P* = 0.005) (Fig. 3C and D). There was no difference compared to parallel measurements of ASL height by conventional fluorescence microscopy in this study (data not shown), and the obtained results correspond well to previous studies using fluorescence microscopy (35, 38). These data show that confocal reflection microscopy is sensitive to detect ASL dysregulation caused by abnormal airway ion transport.

### Confocal reflection microscopy enables ASL height measurement under unperturbed near-physiological conditions

Next, we applied our novel technique to measure the height of unperturbed ASL on primary mouse airway epithelial cultures. ASL height was recorded every 15 minutes for 5 hours and at 24 hours on primary tracheal cultures from βENaC-Tg mice and wild-type littermate controls without previous addition of fluorescent dye and/or volume (Fig. 4A). In contrast to previous measurements using to the fluorescence protocol, ASL height of βENaC-Tg cultures was significantly reduced compared to wild-type cultures at baseline (Fig. 4A and B). Primary tracheal cultures from wild-type mice maintained ASL height at 8.8 ± 0.6 µm whereas cultures from βENaC-Tg mice had a lower ASL height of 5.6 ± 0.3 µm over the course of 5 hours (*P* < 0.001; Fig. 4B). At 24 hours, ASL determined by confocal reflection microscopy was significantly higher than the values obtained with conventional fluorescence microscopy in both wild-type and βENaC-Tg cultures (both *P* < 0.001) (Fig. 4C). These results demonstrate that confocal reflection microscopy under unperturbed conditions enables detection of ASL dysregulation at baseline and indicate that apical addition of dye and/or volume alters 24 hour steady state ASL height determined by fluorescence confocal microscopy.

**Figure 4.**
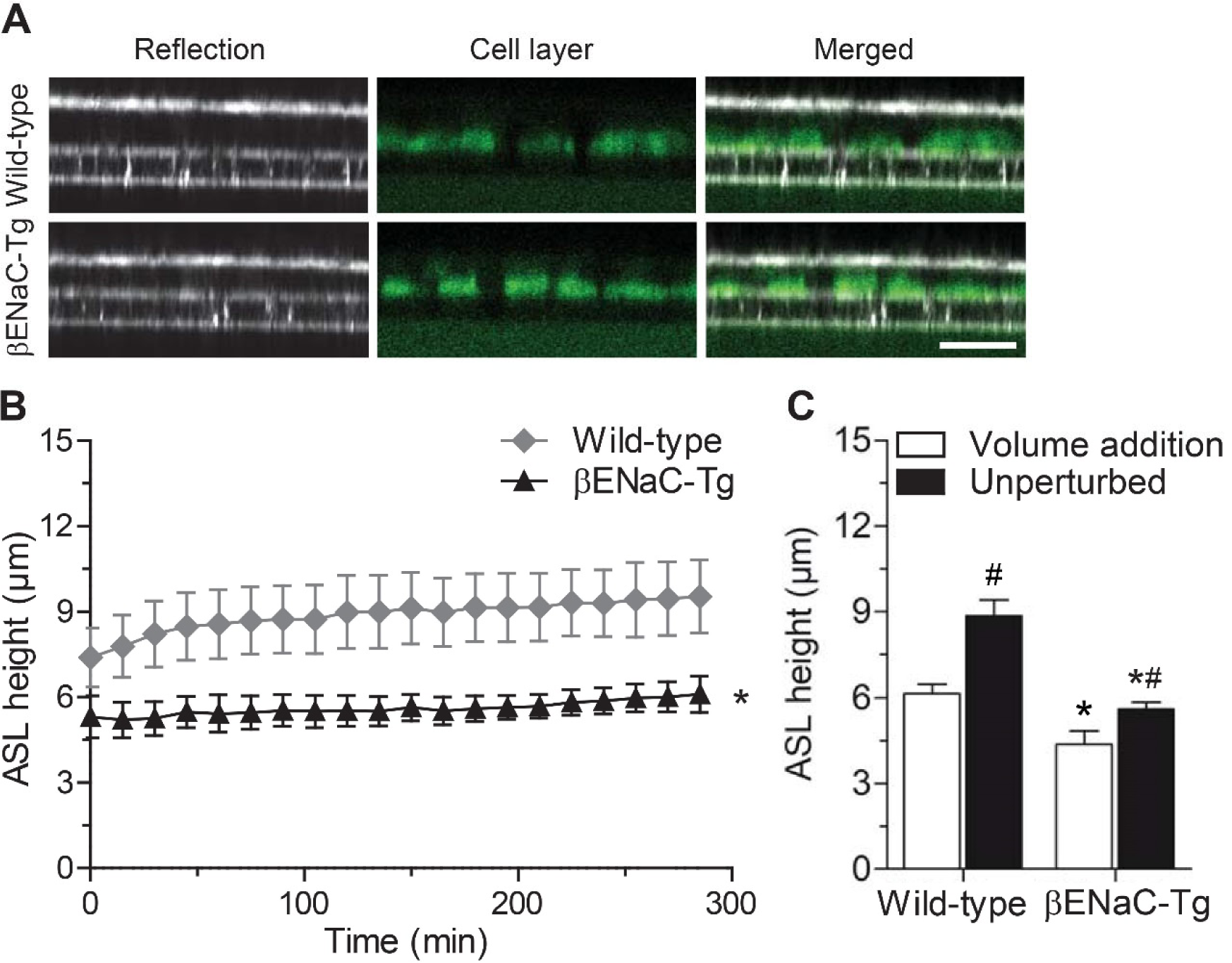
Reflection confocal microscopy detects ASL depletion on unperturbed primary airway cultures from βENaC-Tg mice without apical volume challenge. Primary tracheal epithelial cell cultures from βENaC-Tg and wild-type mice were grown at air-liquid interface for 14 days and ASL height was measured in unperturbed cultures without apical volume challenge. A, B: Representative confocal images of the reflection signals recorded with the 488 nm laser by XZ-scanning and calcein-AM stained cell layer (A) and summary of ASL height measured every 15 minutes over a period of 5 hours (B) on βENaC-Tg vs. wild-type airway cultures without addition of fluorescent dye from the apical side. Scale bar: 30μm. n = 5 cultures per group, *P < 0.01 compared to wild-type. C: Comparison of steady state ASL height determined at 24 hours after the first measurement by reflection confocal microscopy in unperturbed airway cultures vs fluorescence confocal microscopy after apical addition of rhodamine dextran (conditions as in Fig. 3). n = 4-5 cultures per group, \**P* < 0.01 compared to wild-type; ^#^*P* < 0.001 compared to measurements with volume addition.

### Confocal reflection microscopy is sensitive to detect response to pharmacological modulation of ion transport in normal and CF airway epithelia

Finally, we assessed the capability of confocal reflection microscopy to detect abnormalities in ASL height and effects of pharmacological modulation of ion transport in CF vs. non-CF primary airway epithelial cultures under unperturbed near-physiological conditions. At baseline, ASL height was significantly reduced in cultures from CF patients compared to non-CF controls (8.4 ± 0.5 µm vs. 13.5 ± 0.8 µm, *P* < 0.001; Fig. 5A). To study effects of ion transport modulation on ASL height in primary non-CF vs CF cultures, several pharmacological agents acting on fundamental ion transport processes were added to the basolateral bath to avoid perturbation of the ASL. Blocking ENaC-mediated Na^+^ absorption with benzamil resulted in a significant increase of ASL height to 16.3 ± 1.5 µm (*P* = 0.005) in non-CF cultures but had no effect on depleted ASL in CF cultures (Fig. 5A). cAMP-mediated stimulation of Cl^-^ secretion with IBMX and forskolin had no effect on ASL height in either non-CF or CF cultures. Inhibition of the Na^+^-2Cl^−^-K^+^-cotransporter with bumetanide resulted in a significant reduction of ASL height in non-CF cultures to 14.9 ± 1.1 µm (*P* = 0.002), but had no effect on CF cultures (Fig. 5A).

**Figure 5.**
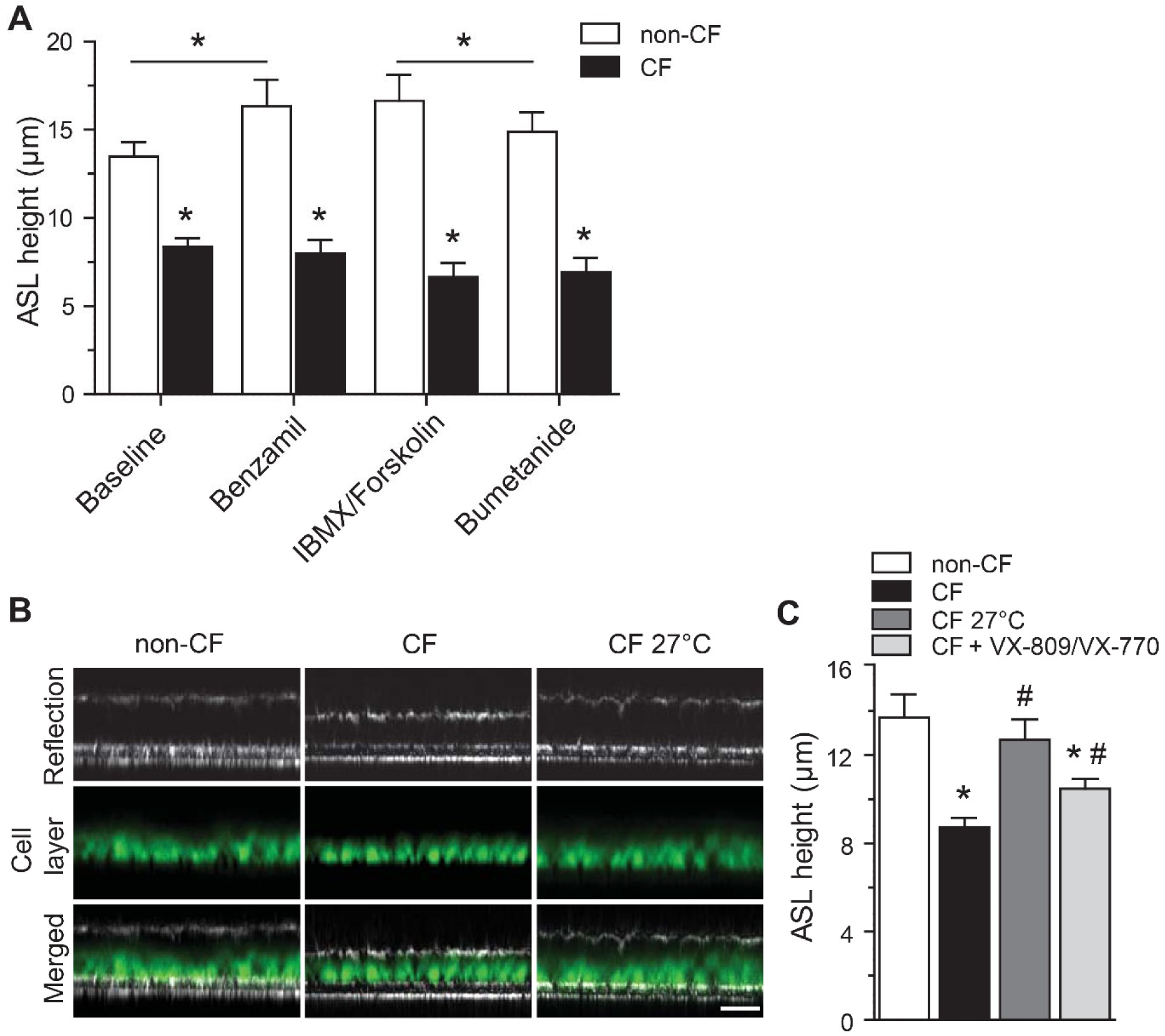
Reflection confocal microscopy detects ASL dysregulation and response to pharmacological modulation of ion transport in non-CF and CF primary airway epithelial cultures. Primary airway epithelial cultures from non-CF controls and CF patients with at least one F508del mutation (Table 1) were grown at air-liquid interface for 14 days and effects of pharmacological modulation of airway ion transport on ASL height was measured by reflection confocal microscopy without apical addition of fluorescent dye. A: Summary of acute effects of inhibition of ENaC by benzamil, cAMP-dependent activation with IBMX and forskolin, and inhibition of transepithelial chloride transport by bumetanide on ASL height on non-CF and CF primary airway epithelial cultures determined by confocal reflection microscopy during sequential addition of compounds to the basolateral compartment. n = 4-6 cultures per group, *P < 0.01 compared to non-CF cultures. B, C: Effects of low temperature (27°C) and treatment with the CFTR modulator combination VX-809/VX-770 (lumacaftor/ivacaftor) on ASL height on F508del-expressing CF primary airway epithelial cultures determined by reflection confocal microscopy. B: Representative confocal images of the reflection signals recorded with the 488 nm laser by XZ-scanning showing steady state ASL height and the calcein-AM stained cell layer. Scale bar: 30μm. C: Summary of ASL height in untreated non-CF vs. CF cultures and effects of incubation at 27°C and treatment with VX-809 and VX-770. n = 5-7 cultures per group, *P < 0.001 compared to non-CF cultures; ^#^P < 0.01 compared to untreated CF cultures.

To study whether confocal reflection microscopy is sensitive to detect changes in ASL height in relation to restoration of CFTR function, primary airway epithelial cell cultures from CF patients with at least one F508del allele (Table 1) were either low temperature corrected by incubation at 27°C (36) or treated with the CFTR corrector VX-809 (lumacaftor) in combination with the potentiator VX-770 (ivacaftor) to improve biosynthesis and function of F508del-CFTR (Fig. 5B,C). ASL height in CF cultures was decreased to ̴60% of non-CF controls (*P* < 0.001). Low temperature correction significantly increased ASL height in CF cultures (*P* = 0.003) to about 90% of the ASL height measured in non-CF cultures (Fig. 5B and C). Treatment with VX-809/VX-770 (lumacaftor/ivacaftor) also improved ASL height on CF cultures to ̴70% (*P* = 0.012; Fig 5C). Collectively, these results demonstrate that confocal reflection microscopy is sensitive to detect response to therapeutic interventions targeting epithelial ion transport including CFTR-mediated Cl^-^ secretion at the level of ASL height.

## DISCUSSION

In this study, we present confocal reflection microscopy as a new method that enables quantitative assessment of ASL on highly-differentiated primary airway epithelial cultures under unperturbed near-physiological conditions by detection of refracted light (Fig. 1). First, we validated the newly developed method by side-by-side comparison to conventional ASL height measurements requiring fluorescent dye addition using primary mouse airway epithelial cell cultures at ALI (Fig. 2 and 3). Second, we show that confocal reflection microscopy is sensitive to detect the effects of Na^+^ hyperabsorption on ASL height in primary mouse airway epithelial cell cultures (Fig. 4). Finally, we demonstrate that this method is sensitive to detect effects of acute as well as chronic pharmacological modulation of ion transport properties on ASL height in CF and non-CF primary human airway epithelial cell cultures at ALI (Fig. 5).

In the past, in addition to conventional confocal fluorescence microscopy, measurements of ASL regulation were also performed using optical coherence tomography (OCT) or synchrotron techniques (41, 42). While these techniques provided valuable insights in the dynamic processes governing airway surface hydration and mucociliary clearance, they have several limitations: OCT offers high-resolution imaging but is limited in its ability to measure ASL thickness accurately due to refractive index mismatch and boundary effects (43). Similarly, synchrotron techniques provide detailed information but are constrained by accessibility and complexity (42, 44). The development of confocal reflection microscopy represents a significant advancement in ASL measurement techniques. Unlike the previously used methods, confocal reflection microscopy by a widely available standard confocal microscope offers non-invasive and quantitative assessment of ASL height under unperturbed conditions. By detecting refracted light, this method circumvents the need for exogenous dyes or volume manipulations, providing a more accurate representation of ASL regulation under near-physiological conditions (Fig. 1).

In our study, we compared ASL measurements obtained by confocal reflection microscopy with those obtained using conventional confocal fluorescence microscopy (Fig. 2 and 3). Measurements with confocal reflection microscopy showed that unperturbed ASL in ALI cultures from wild-type mice is higher than the ASL measured 24 hours after volume addition as determined by conventional confocal fluorescence microscopy (Fig. 4). This difference suggests that the perturbations induced by the addition of extra liquid may have prolonged effects on volume regulation, potentially taking a longer time to rebalance soluble endogenous regulators of ASL and return to equilibrium than previously assumed (45-48). Several mechanisms involved in sensing surface hydration could contribute to these observed differences. First, volume addition has been reported to activate near-silent ENaC channels by dilution of endogenous inhibitors of the channel activating proteases (CAP), such as CAP1/prostasin (49, 50). Additionally, prolonged volume changes may impact the transcriptional regulation of proteins involved in ion transport, such as increased expression of the endogenous serine protease prostasin (45). Thus, our data suggest that a prolonged increase in ENaC activity following volume challenge may alter ASL height when using conventional confocal fluorescence microscopy compared to confocal reflection microscopy that does not require a volume challenge of the cultures. Along these lines, our data demonstrate that there is no response to the ENaC blocker benzamil in human CF primary airway epithelial cells under unperturbed thin film conditions (Fig. 5) most likely due to near silent ENaC channels under physiological conditions, where cleavage-activation by proteases including CAPs is prevented by endogenous antiproteases (51). In comparison, in non-CF cultures with an initial ASL height almost twice as high as in CF cultures, endogenous protease inhibitors may be diluted resulting in active ENaC channels, therefore ASL is increased after inhibition of ENaC-mediated absorption by benzamil. Subsequent stimulation of CFTR-mediated Cl^-^ secretion did not result in further increase of ASL height in airway epithelial cells from non-CF subjects (Fig. 5) since benzamil-induced apical membrane hyperpolarization may have already stimulated fluid secretion (52, 53). Subsequent inhibition of the Na^+^-K^+^-2Cl^-^ co-transporter, resulting in a reduced driving force for Cl^-^ secretion through reduced Cl^-^ uptake and drop in intracellular Cl^-^ concentration lead to a net decrease in in ASL height in non-CF cultures (Fig. 5).

In addition to studies of ASL regulation on non-CF airway cultures, we also assessed the effect of pharmacological rescue of CFTR by the combination of the corrector VX-809 (lumacaftor) and the potentiator VX-770 (ivacaftor) in nasal epithelial cultures from patients with CF homozygous for the F508del mutation. We show for the first time that treatment of human airway epithelial cells of CF patients with VX-809 and VX-770 partially restores ASL height by about 30% of normal under unperturbed steady state conditions (Fig. 5). These results are consistent with previous studies by van Goor and colleagues showing that chronic exposure with VX-809 followed by acute addition of VX-770 restored CFTR-mediated chloride transport across human bronchial epithelial cultures from F508del homozygous patients to approximately 25% of that measured in non-CF human bronchial epithelial cells and that this increase in Cl^-^ transport translated into an increase in the ASL height (14). Meanwhile this level of functional rescue was confirmed in observational studies in F508del homozygous CF patients where effects of lumacaftor-ivacaftor therapy on *in vivo* CFTR function was determined by nasal potential difference and intestinal current measurements and was found to improve CFTR function in the airway and intestinal epithelium to levels of 10 to 30% of normal CFTR activity (54, 55).

Collectively, our findings underscore the complexity of ASL regulation and support the use of confocal reflection microscopy for quantitative studies of the unperturbed thin ASL covering airway surfaces in health and disease. Future applications of this technique may include studies of the mechanisms underlying ASL (dys)regulation as well as therapeutic approaches to improve ASL volume and mucociliary clearance. For example, in CF ̴10% of patients still have a high unmet need as they are not eligible for CFTR modulators. In these patients, a personalized medicine approach including confocal reflection microscopy may help to determine response to therapy and provide access to approved CFTR modulator therapies (13, 56-58). Beyond CF, studies on ASL dysregulation may provide insights into the pathophysiology of other muco-obstructive lung diseases such as COPD and non-CF bronchiectasis (8, 59). In this context, there is interest in targeting alternative ion channels such as ENaC and the alternative Cl^-^ channel TMEM16A to increase ASL volume and improve mucociliary clearance in these muco-obstructive lung diseases (8, 20, 59-61). Quantitative assessment of effects of therapeutic approaches on ASL height on airway epithelial cultures under unperturbed conditions by confocal reflection microscopy may facilitate the discovery and preclinical development of novel therapies aiming to improve airway surface hydration and mucus clearance.

In summary, our study highlights the importance of considering the intricate mechanisms underlying the regulation of epithelial ion transport as well as regulation of the thin ASL covering airway surfaces for studies of the pathophysiology and the development of novel therapeutic strategies for muco-obstructive lung diseases. Our data support that confocal reflection microscopy may serve as a valuable tool for elucidating disease mechanisms and guiding the development of novel therapies aiming at restoring ASL homeostasis and mucus clearance in CF and potentially other muco-obstructive lung diseases.

## DATA AVAILABILITY

Data will be made available upon reasonable request.

## ACKNOWLEDGMENTS

We thank J. Schatterny and S. Butz for excellent technical assistance.

## AUTHOR CONTRIBUTIONS

Conception and design: A.S.A, M.L., M.A.M., J.D. Acquisition, analysis and interpretation of data: A.S.A, M.L., H.S., T.A., S.Y.G., A.B., I.B., S.B., R.P., M.A.M., J.D. Writing the manuscript or revising it critically for important intellectual content: A.S.A, M.L., H.S., T.A., S.Y.G., A.B., I.B., S.B., R.P., M.A.M., J.D.

